# Breaking barriers: the fungal toxin candidalysin disrupts epithelial integrity and induces inflammation in a gut-on-chip model

**DOI:** 10.1101/2024.10.30.621017

**Authors:** Moran Morelli, Karla Queiroz

## Abstract

*Candida albicans* is an opportunistic pathogenic yeast commonly found in the gastrointestinal tract, vagina, and mouth of healthy humans. Under certain conditions, it can become invasive, causing mucosal or life-threatening systemic infections. One mechanism used by *C*.*albicans* to breach the epithelial barrier is the secretion of candidalysin, a cytolytic peptide toxin. Candidalysin damages epithelial membranes and activates the innate epithelial immune response, making it key to *C*.*albicans’* pathogenicity and a promising therapeutic target. Although candidalysin mediates *C. albicans* translocation through intestinal layers, its impact on epithelial responses is not fully understood.

This study aims to characterize this response and develop scalable, quantitative methodologies to assess candidalysin’s toxicological effects using gut-on-chip models.

We used the OrganoPlate®, a microfluidic platform to culture up to 64 perfused, membrane-free intestinal epithelial tubes. We exposed Caco-2 tubes to candidalysin and evaluated their response with trans-epithelial electrical resistance (TEER), protein detection, and immunostaining. We then validated our findings in a proof-of-concept experiment using human intestinal organoid tubules.

Candidalysin impaired barrier integrity, as indicated by decreased TEER and increased permeability in a fluorescent dye assay. It also induced actin remodeling and DRAQ7™ dye uptake, a marker of cell permeability. This disruption was associated with the release of LDH, cytokines, and the antimicrobial peptide LL37, suggesting cellular damage, inflammation, and antimicrobial activity.

This study strengthens our understanding of candidalysin’s role in *C. albicans* pathogenesis and suggests new therapeutic strategies targeting this toxin. Moreover, the use of patient-derived organoids shows promise for capturing patient heterogeneity and developing personalized treatments.

**Graphical Abstract:** 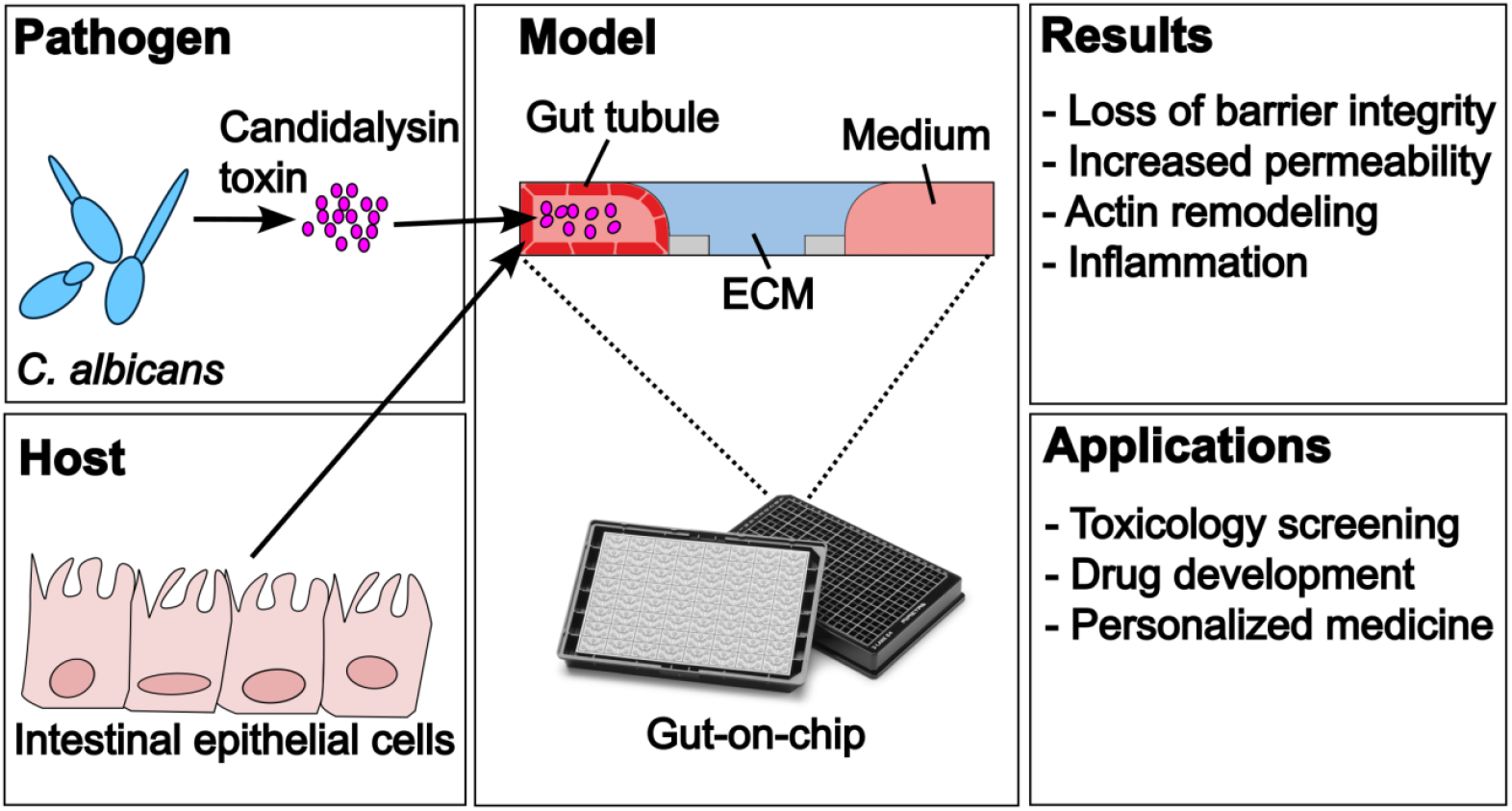

## 1 Introduction

The intestinal mucosa is the first line of defense against microbial invasion. Intestinal epithelial cells (IECs) form a barrier, which separates the sterile host environment from the gut microbiota. Perturbations of the host-microbial interplay such as an imbalance in the microbial community, a breach in the barrier, or an impairment of the host’s immune system can lead to disease[1].

Candida species are among the leading fungal pathogens, significantly contributing to morbidity and mortality[2]. Among these, *C. albicans* remains the most prevalent cause of life-threatening systemic candidiasis and is a major contributor to nosocomial infections, particularly in Intensive Care Units (ICUs)[2, 3]. *C. albicans* is an opportunistic pathogen that frequently inhabits the mucosal surfaces of humans, with its main reservoir being the gastrointestinal tract[4–7]. A compromised immune system and the use of broad-spectrum antibiotics can induce the transition of *C. albicans* from commensalism to pathogenicity, where it can breach the intestinal barrier and reach the bloodstream to cause systemic candidiasis[8–10]. One of the key mechanisms mediating the translocation of *C. albicans* through the intestinal layers is the release of the virulence factor candidalysin[11, 12].

Candidalysin is a cytolytic peptide playing a dual role in *C. albicans* infection. On one hand, it acts as a virulence factor by directly damaging the host’s epithelium through the formation of membrane pores[11, 13, 14]. On the other hand, candidalysin acts as an immunomodulatory molecule recognized by the host to initiate an immune response through p38 and EGFR-ERK signaling[11, 15, 16]. Characterizing the epithelial response to candidalysin is therefore of critical importance to better manage candidiasis.

So far, most of the *in vitro* research exploring the mechanisms of candidalysin has been done in oral epithelial cell lines, where candidalysin has been shown to trigger the release of cytokines, chemokines, alarmins, and antimicrobial peptides[16–19]; to induce epithelial stress such as mitochondrial dysfunction, ATP depletion, and epithelial necrosis[20]; and to trigger Ca^2+^ influx and breakdown of F-actin[21]. Nevertheless, although it is known that the main reservoir of *C. albicans* is the gut[4, 22], not many studies explored the effect of candidalysin on intestinal epithelial cells (IECs).

Allert and colleagues investigated the translocation process of *C. albicans* in IECs *in vitro* by screening for >2000 mutants with deleted genes coding for translocation and cellular damage. They showed that the mutant without candidalysin displayed moderate loss of barrier integrity, caused almost no damage, and exhibited a reduced translocation in the Caco-2 brush border subclone C2BBe1, indicating that the toxin is essential for translocation of the fungus through the intestinal wall[10].

A follow up study investigated the mechanisms of *C. albicans* translocation across the intestinal epithelial barrier focusing on fungal and host activities involved in the process. They found that intestinal translocation was promoted by acquisition of host-cell zinc by *C. albicans* while at the same time host NFκB signaling was protecting epithelial integrity, mitigating candidalysin-induced damage and limiting fungal translocation[23].

While previous studies have explored the role of candidalysin in epithelial damage and translocation, they primarily utilized static models or non-gut epithelial cells, which are limited in their ability to dynamically assess barrier function and immune responses within a physiologically relevant gut environment. Additionally, these models often lack scalable and quantitative methodologies, restricting their utility for high-throughput screening and therapeutic development.

Here, we used a membrane-free gut-on-chip model to evaluate the intestinal response to candidalysin in the cell line Caco-2, using scalable and quantitative readouts. We characterized the response to the toxin by studying the barrier integrity, morphology, permeability, and release of epithelial mediators at the luminal side of the model. To validate our findings in a more relevant gut-on-chip model, we included a proof-of-concept experiment using patient-derived colon organoids to study barrier integrity, morphology, and permeability after candidalysin exposure.

In summary, our study presents a novel gut-on-chip platform and robust, scalable readouts to evaluate candidalysin’s effects on intestinal epithelial cells. This system is ideal for phenotypic screenings to identify therapeutic molecules and can evolve to include other relevant cell types, enabling comprehensive, physiologically relevant models for both toxicological and for personalized drug screening and the identification of drugs targeting microbial virulence factors.

## 2 Materials and Methods

### 2.1 Cell Culture

#### 2.1.1 OrganoReady Colon Caco-2

OrganoReady Colon Caco-2 3-lane 40 (MI-OR-CC-01, MIMETAS B.V.) were cultured according to the manufacturer’s instructions (**Fig. 1**). Medium was replaced with Caco-2 medium on the day of receiving. OrganoReady Colon Caco-2 3-lane 40 are ready to use Caco-2 tubes in OrganoPlate that follow a similar process than what was described by Trietsch and colleagues[24]. Perfusion flow was maintained by placing the plate on OrganoFlow® rocker (MIMETAS B.V., MI-OFPR-L) set at 7 degrees with 8-minute intervals optimized for the 3-lane 40. On the second day after receiving (day 6 after seeding), Caco-2 medium was refreshed. Exposures were performed on day 4 after receiving (day 8 after seeding).

**Fig. 1.**
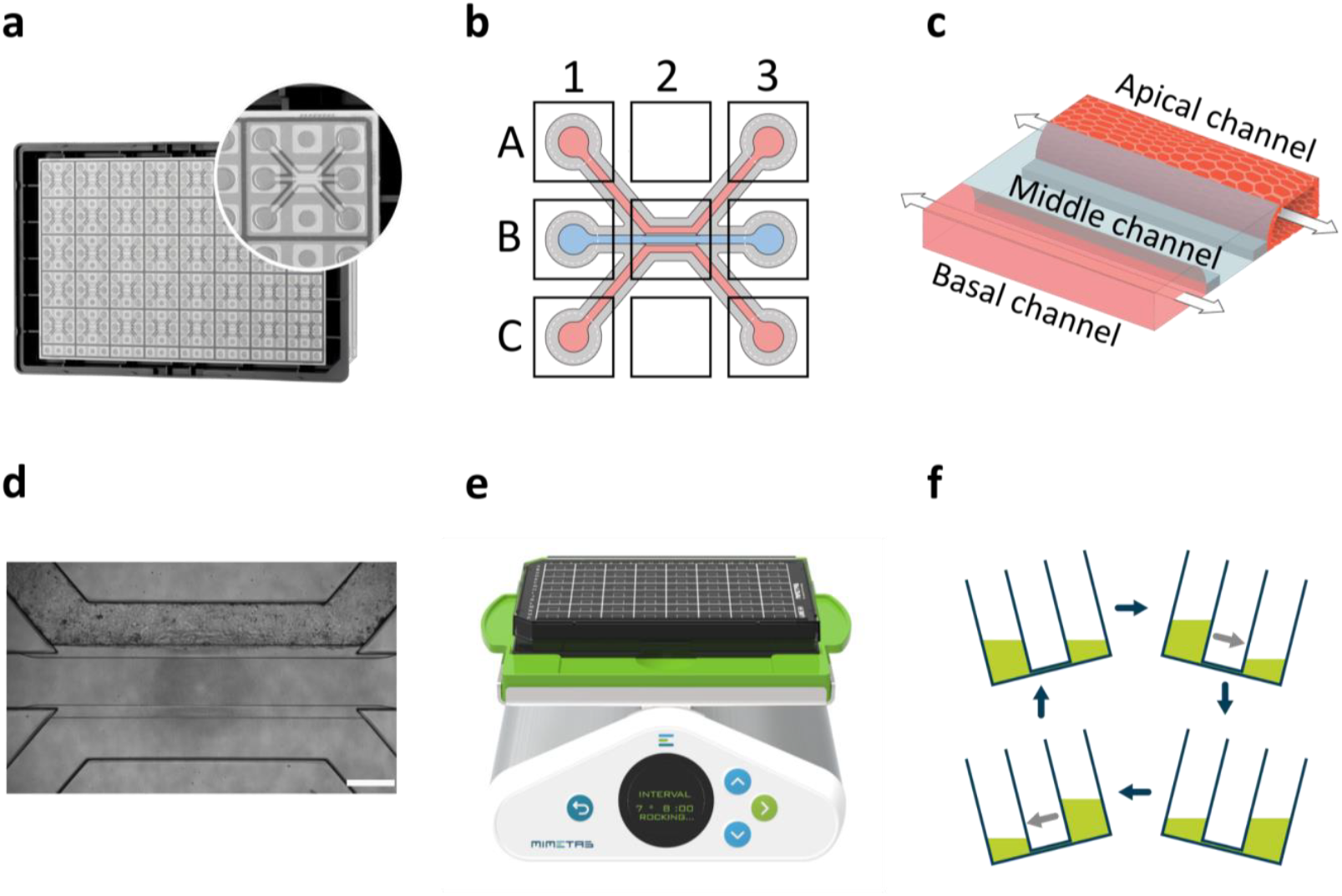
OrganoReady Caco-2 model. **(a)** Picture of an OrganoPlate 3-lane 40 with one microfluidic chip enlarged. (**b**) Illustration of a microfluidic chip. Apical channel can be accessed through A1 and A3. Middle channel can be accessed through B1 and B3. Basal channel can be accessed through C1 and C3. (**c**) Schematic of a Caco-2 tubule (apical channel) against a collagen ECM (middle channel). Basal channel contains culture medium. Arrows indicate flow direction. (**d**) Phase-contrast image of a Caco-2 tubule in the OrganoPLate 3-lane 40, scale bar is 100 μM. (**e**) OrganoPlate on an OrganoFlow rocker. The rocker angle is set at 7 degrees and oscillates every 8 minutes. (**f**) Illustration of bi-directional flow in the OrganoPlate

#### 2.1.2 OrganoReady Colon Organoid

OrganoReady Colon Organoid plates (OrganoReady Colon Organoid 3-lane 64, MI-OR-CORG-02, MIMETAS B.V.) were used to study the effect of candidalysin in colon organoids. Each plate contained 64 chips with colon organoid tubules, cultured using the following media: OrganoMedium Colon Organoid-ARM (Apical Recovery Medium), BRM (Basolateral Recovery Medium), ACM (Apical Culture Medium), and BCM (Basolateral Culture Medium), per the manufacturer’s instructions. Perfusion flow was maintained on an OrganoFlow rocker (MIMETAS B.V., MI-OFPR-L) set at 14 degrees with 8-minute intervals optimized for the 3-lane 64. The tubules were derived from the healthy portion of a sigmoid biopsy from a 58-year-old Caucasian female with colorectal cancer.

### 2.2 Candidalysin Exposure

Candidalysin (SIIGIIMGILGNIPQVIQIIMSIVKAFKGNK) was synthesized by Peptide Protein Research Ltd (UK) and reconstituted in sterile purified water to 5 mg/mL (1.5 mM) for storage at -20°C, prior to further dilution for individual experiments.

For exposures in Caco-2 tubules, medium was replaced with serum-free medium in all channels 24h prior to exposure. Then, the medium of the apical channel was replaced with serum-free medium containing candidalysin at the specified concentration.

Due to their increased sensitivity and the limitation of modifying their culture medium, colon organoids tubules were not serum starved. Instead, all channels were washed with HBSS (Hank’s Balanced Salt Solution) prior to adding HBSS containing the specified concentration of candidalysin in the apical channel.

### 2.3 Protein Detection

Supernatant were collected from the apical channel and centrifuged for 10 min at 1500 rpm prior to aliquoting and storage at -80 °C.

LL37 ELISA kits were purchased from Elabscience (UK) and performed according to the manufacturer’s instructions. Absorbance was measured using a Spark Cyto plate reader (TECAN, Switzerland).

G-CSF, GM-CSF, IL-1 alpha, IL-1 beta, IL-6, IL-8, IP-10, MCP-1 (CCL2), MIP-3 alpha (CCL20), S100A8/A9 were measured using a Luminex MAGPIX instrument (Thermo, USA) and a custom Procartaplex assay kit (Thermo, USA), following the manufacturer’s instructions. Plates were measured using a Luminex MAGPIX® instrument and the Luminex xPONENT® software (version 4.3). ProcartaPlex Analysis App was used to determine analyte concentrations.

### 2.4 Trans-Epithelial-Electrical-Resistance (TEER)

Trans-epithelial electrical resistance (TEER) was measured using an automated multichannel impedance spectrometer (OrganoTEER®, **Fig. 2 a-c**) to evaluate the integrity of the gut barrier[25]. An electrode board, designed to fit the 3-lane OrganoPlate was sterilized with 70% ethanol at least 1 hour before measurement. The OrganoPlate was taken out of the incubator and equilibrated at room temperature for 30 min prior to the measurement to eliminate any temperature or flow effect. Baseline measurement was performed right after equilibration, then TEER was further measured at the indicated timepoints after candidalysin exposure.

**Fig. 2.**
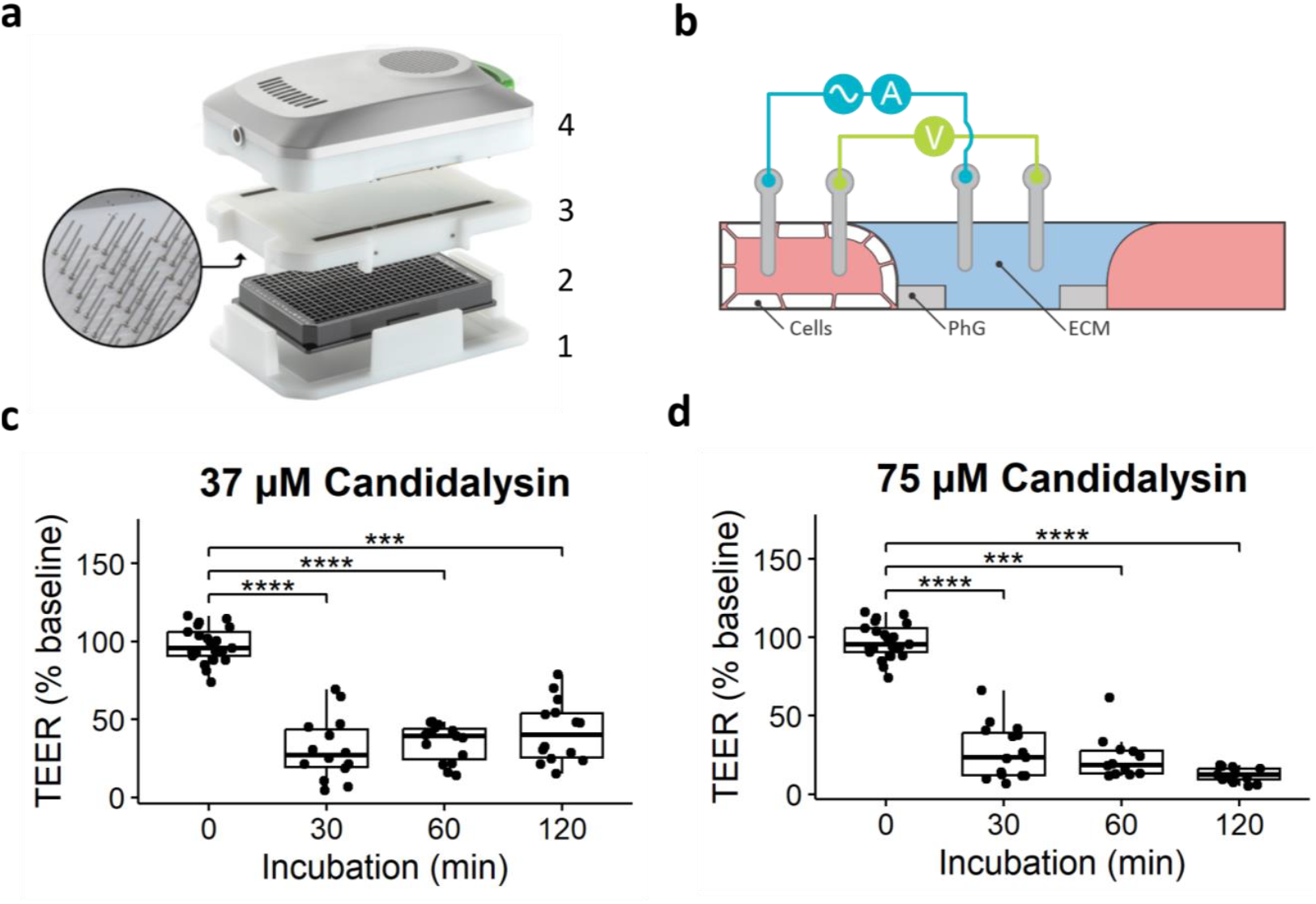
TEER measurements of Caco-2 tubules exposed to candidalysin. (**a**) Photograph of the OrganoTEER setup: the plate holder (1) holds the OrganoPlate (2) in which the electrode board (3) is positioned and connected to the measuring device (4). (**b**) Illustration of a cross-section of a microfluidic chip and positioning of the electrodes. PhG: phaseguide technology, allowing membrane-free separation of the channels. ECM: extracellular matrix. (**c**) TEER measurements after 30min, 60min, and 120min exposure to 37 μM candidalysin, and (**d**) 75 μM candidalysin. Results are expressed as boxplot with individual measurements for each chip. Each chip is normalized to its baseline value before exposure. Data was analyzed using Kruskal Wallis test with Dunn’s post-hoc test (*p ≤ 0.05; **p ≤ 0.01; ***p ≤ 0.001, ****p ≤ 0.0001, N=3, n=3-8)

For Caco-2 the OrganoTEER was set on “high TEER” and we used a TEER threshold of 350 Ω/cm^2^ at baseline, meaning that all tubules below 350 Ω/cm^2^ before exposure were excluded from the analysis. For colon organoids the OrganoTEER was set on “medium TEER” and we used a TEER threshold of 100 Ω/cm^2^ at baseline, meaning that all tubules below 100 Ω/cm^2^ before exposure were excluded from the analysis.

### 2.5 Barrier Integrity Assay

To further investigate the barrier integrity, leakage of fluorescent molecules from the lumen to the ECM compartment was evaluated after exposure (**Fig. 3 a-c**). First, medium from all inlet and outlets was removed. Then 25 μL serum-free medium was added to the inlet and outlets of the middle and basal channel. Next, a dye solution consisting of Sodium fluorescein (0.4 kDa, 10 μg/ml, Sigma-Aldrich, USA) and TRITC-dextran (4.4 kDa, 500 μg/ml, Sigma-Aldrich, USA) in serum-free medium was added to the inlet and outlet of the apical channel (40 and 30 μL, respectively). Then, the OrganoPlate was placed in an ImageXpress XLS Micro (Molecular Devices, USA) and imaged every 2 minutes for 7 timepoints. Leakage from the lumen into the ECM compartment was quantified according to the protocol described by Soragni et al.[26]

**Fig. 3.**
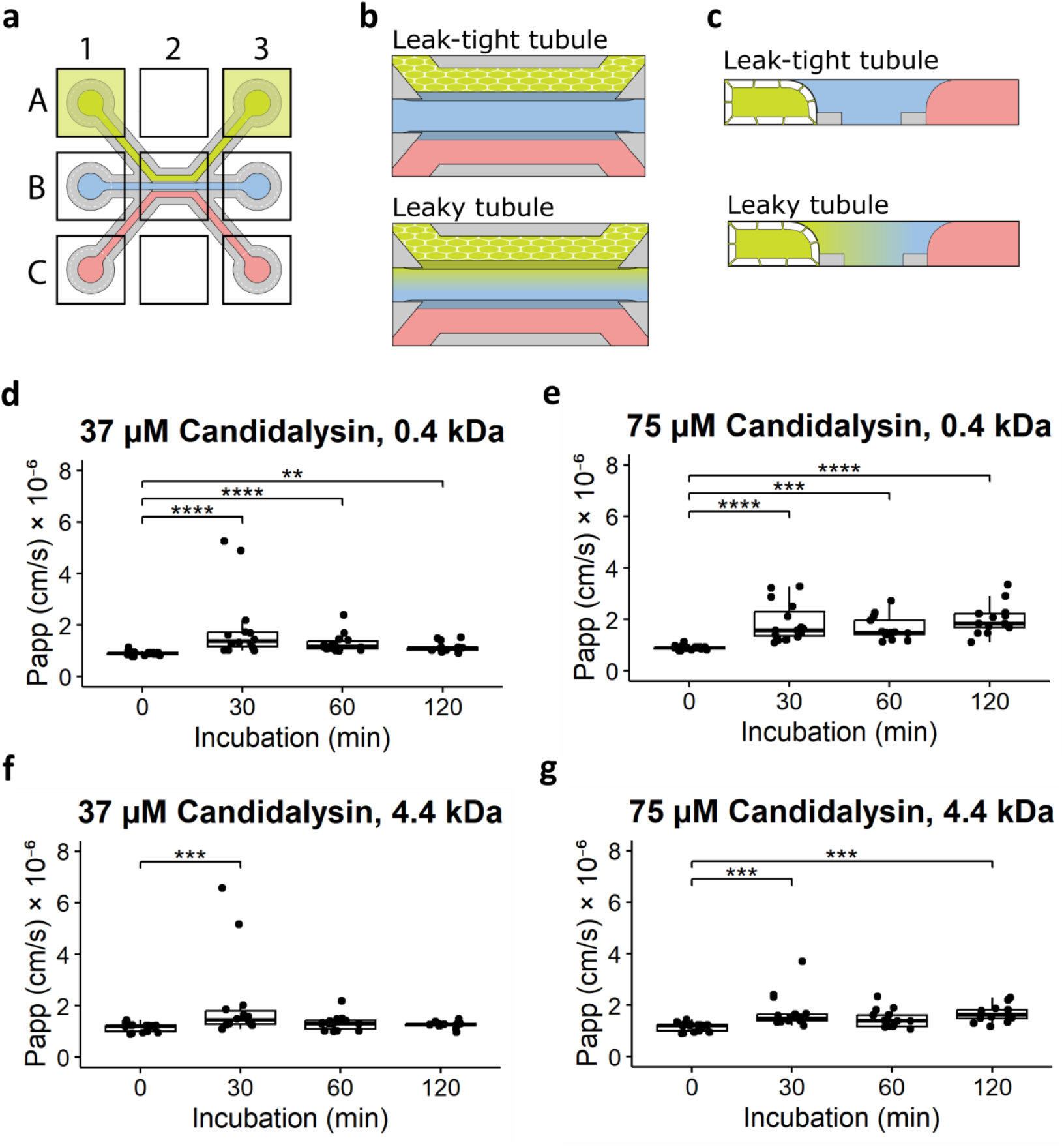
Caco-2 tubule permeability to fluorescent dyes after candidalysin exposure. (**a**) Illustration of a chip with the dye added to the lumen of the Caco-2 tubule. (**b**) Enlarged view of a chip with dye perfused in the lumen of the tubule. Top illustration shows a leak-tight tubule; bottom illustration shows a leaky tubule. (**c**) Cross-section of a chip to show a leak-tight tubule at the top and a leaky tubule at the bottom. (**d**) Permeability of Caco-2 tubule to 0.4 kDa dye after 30, 60, and 120min exposure to 37 μM, and (**e**) 75 μM candidalysin. (**f**) Permeability of Caco-2 tubule to 4.4 kDa dye after 30, 60, and 120min exposure to 37 μM, and (**g**) 75 μM candidalysin. (Results are expressed as boxplot with individual measurements for each chip. Data was analyzed using Kruskal Wallis test with Dunn’s post-hoc test (*p ≤ 0.05; **p ≤ 0.01; ***p ≤ 0.001, ****p ≤ 0.0001, N=3, n=3-7)

### 2.6 Cell Damage (LDH) Assay

Lactate dehydrogenase (LDH) activity was measured using the LDH-Glo™ Cytotoxicity Assay kit (Promega, USA), following the manufacturer’s instructions. Supernatant were collected from the lumen and centrifuged for 10 min at 1500 rpm, prior to dilution in LDH assay buffer (1:100) and storage at -80 °C. Luminescence was measured in 96W half area white microplates (Greiner bio-one, Austria) using a Spark Cyto plate reader (TECAN, Switzerland).

### 2.7 Immunohistochemistry

For visual characterization of the effect of candidalysin on actin cytoskeleton and cell permeability, tubules were directly fixed and stained on the OrganoPlate based on the protocol described by Trietsch et al.[24]

#### 2.7.1 DRAQ7 staining

Before fixation, medium in the tubule inlets and outlets was replaced by 20μL of DRAQ7 dye (Biostatus, DR71000) diluted 1:100 in serum-free EMEM (Caco-2) or HBSS (colon organoids), and cells were incubated during 30 min under continuous perfusion inside the incubator. OrganoPlates were then fixed and stained following the previously explained protocol.

#### 2.7.2 Fixation

Intestinal tubules were fixed with 3.7% formaldehyde (Sigma, 252,549) in HBSS with Calcium and Magnesium (Thermo Scientific, 14,025,092) for 15 min, washed twice with phosphate-buffered saline (PBS; Gibco, 70,013,065) for 5 min and then stored with 50μL PBS per well at 4 °C until further staining.

#### 2.7.3 Actin and Nuclear Staining

Intestinal tubules cells were permeabilized with 0.03% Triton X-100 (Sigma, T8787) in PBS for 10 min and washed twice with 4% FBS in PBS solution. Actin and nuclear stainings were performed using the direct stains ActinGreen™ 488 ReadyProbes™ Reagent (Invitrogen, R37110) and NucBlue™ Fixed Cell ReadyProbes Reagent (Hoechst 33342, Invitrogen, R37605). Direct staining was performed following the manufacturer’s instructions and under constant flow.

For actin and Hoechst staining, two drops/ml were added in PBS and 20μl of the staining solution was added to the tube inlet and outlet (20μL each) and incubated for 30min at room temperature under continuous perfusion. Plates were then washed twice for 5 minutes with PBS and the bottom of the tubules was imaged as max projection using an ImageXpress Micro Confocal microscope (Molecular Devices, USA).

### 2.8 Intestinal tubules visualization and imaging

Tubules were imaged using the ImageXpress® Micro XLS (Molecular Devices) and Micro XLS-C High Content Imaging Systems (Molecular Devices) and processed using Fiji34 to enhance contrast and improve visualization. To monitor the integrity of the tubules, phase-contrast images were recorded before and after exposure to candidalysin. This was routinely performed immediately after measurement of baseline TEER and after the required exposure time. Fixed and stained OrganoPlates were stored at 4 °C until imaging and equilibrated at room temperature at least 30 min before imaging. Maximum intensity projection images were saved as TIFF files after confocal imaging of stained cells.

### 2.9 Staining quantification

Open-source cell image analysis software CellProfiler™ (version 4.2.5) was used to process visual immunofluorescence images. An image analysis pipeline was designed to quantify the number and area of actin clumps and the number of cells, by segmentation of the actin clumps and nuclei and measurement of the total actin clump area. This pipeline was used to process the TIFF files that captured actin staining or nuclei staining, correspondingly. Parallelly, another pipeline was designed to process the TIFF files capturing DRAQ7 and nuclei staining, that allowed segmentation of DRAQ7-positive cells and nuclei, to quantify the percentage of DRAQ7-positive (DRAQ7 +) cells.

### 2.10 Statistics

Data analysis and visualization were conducted using R version 4.2.2 and RStudio version 2023.03.0-386.

All experiments were performed three times in an independent manner (N = 3), and the number of technical replicates per concentration (n) is indicated in the caption of each figure. Normality of data distribution and homogeneity of variances were assessed using the Shapiro-Wilk test and Levene’s test, respectively. Differences among two groups were compared using independent T-test or Wilcoxon Rank-Sum Test (Mann-Whitney U Test). Differences among three groups or more were compared using one-way ANOVA followed by post-hoc Tukey’s HSD, or Kruskal–Wallis tests followed by Dunn’s test, to obtain pairwise comparisons between the effect of exposure to the various candidalysin concentrations and exposure. P values were adjusted for multiple comparisons using Bonferroni method. Adjusted P values of 5% or lower were considered statistically significant and were reported with asterisks: *p ≤ 0.05; **p ≤ 0.01; ***p ≤ 0.001, ****p ≤ 0.0001. P values above 5% were considered not statistically significant (ns) and were not reported.

## 3 Results

### 3.1 Candidalysin impairs barrier integrity of Caco-2 tubules and increases their permeability

To determine candidalysin-mediated effects on the intestinal barrier integrity, we used the OrganoTEER (**Fig. 2 a-b**) to measure TEER of the Caco-2 tubules after 30min, 1h, and 2h candidalysin exposure (37 and 75 μM). Both concentrations significantly decreased TEER after all timepoints compared to vehicle (**Fig. 2 c, d**). When comparing the effect over time, although 37 μM candidalysin exposure showed a trend of recovering TEER over time, and 75 μM candidalysin showed a trend of decreasing TEER over time, these changes were not statistically significant (p > 0.05) (**Figure 2 c, d**).

We next investigated the permeability of the Caco-2 tubules after 37 and 75 μM candidalysin exposure, using fluorescent molecules of two different sizes. Permeability to sodium fluorescein (0.4 kDa) was significantly increased after 30, 60, and 120min incubation with 37 μM candidalysin (**Fig. 3 d**), while permeability to dextran 4.4 kDa was significantly increased only after 30min 37 μM candidalysin exposure (**Fig. 3 f**). 75 μM candidalysin increased permeability to both dyes after all incubation times (**Fig. 3 e, g**). Following evaluation of permeability across various candidalysin exposure conditions, we selected the 2-hour incubation with 75 μM candidalysin for further analysis.

We then assessed whether the expression of the permeability marker DRAQ7 was increased after 75 μM candidalysin exposure for 2h when compared to vehicle. **Fig. 4 a** shows images of representative chips exposed to vehicle or candidalysin and stained for DRAQ7 and Hoechst. Permeable cells’ nuclei and cell count (total nuclei) quantification are shown in middle and bottom rows, respectively. 75 μM candidalysin significantly increased the number of DRAQ7 positive cells compared to vehicle (**Fig. 4 b**).

**Fig. 4.**
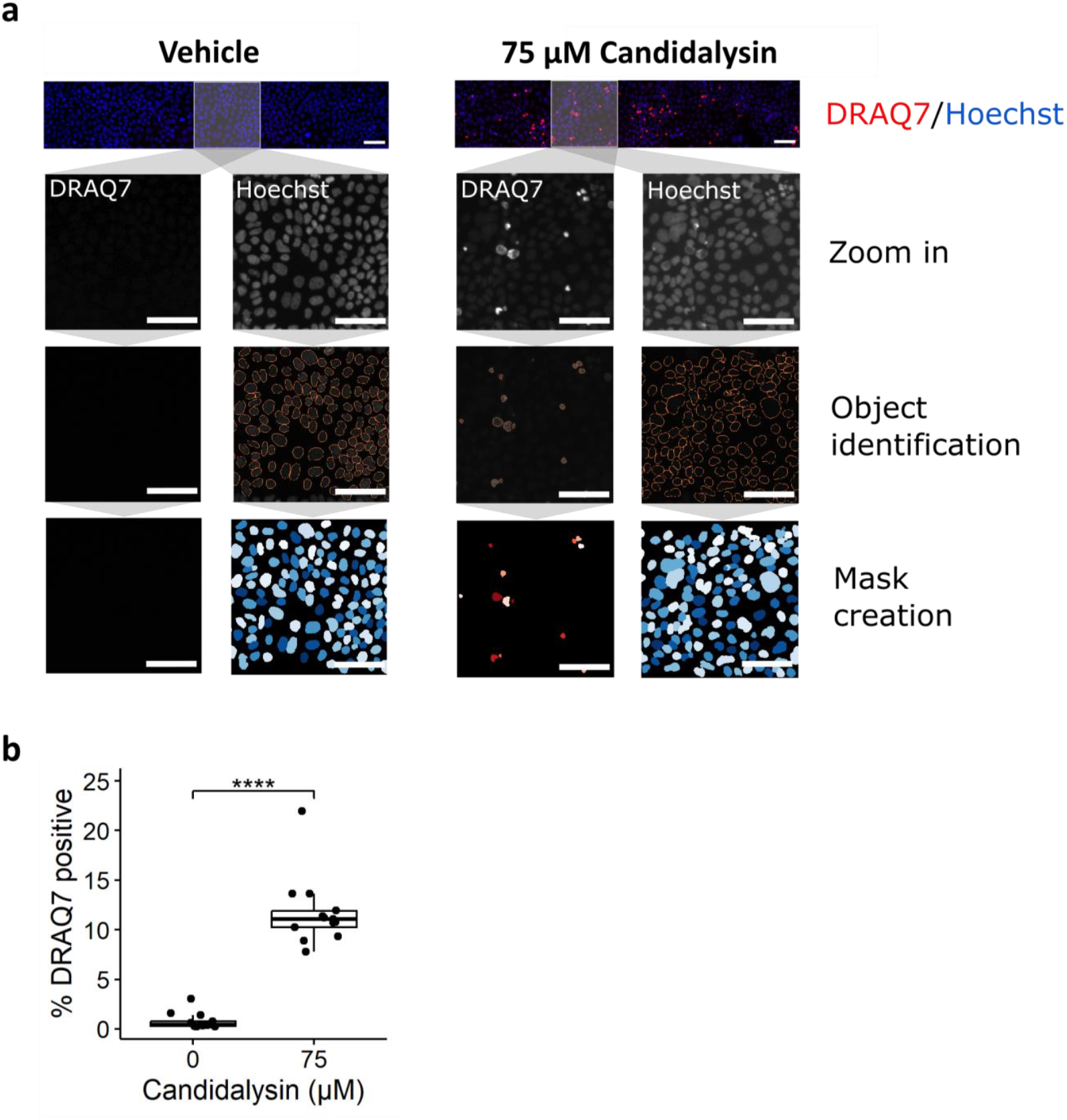
Candidalysin increases permeability of IECs. Caco-2 tubules exposed to candidalysin for 2h were stained with DRAQ7 nuclear dye, which only penetrates permeabilized or dead cells, and with Hoechst. (**a**) A CellProfiler pipeline was designed to identify and quantify DRAQ7-positive and Hoechst-positive nuclei from confocal images. DRAQ7 + objects were filtered to only consider those overlapping with Hoechst positive objects. Pictures are representative of 3 independent experiments (N=3). Scalebar is 100 μM. (**b**) The number of DRAQ7 + nuclei was normalized against the total number of nuclei (Hoechst positive) to give a percentage of DRAQ7 positive cells.) Results are expressed as boxplot with individual measurements for each chip. Data was analyzed using Wilcoxon Rank-Sum Test (****p ≤ 0.0001, N=3, n=4-5)

### 3.2 Candidalysin induces actin remodeling in Caco-2 tubules

Next, we investigated the morphology of the tubules after candidalysin exposure. Cells were exposed to vehicle or 75 μM candidalysin for 2h. **Fig. 5 a** shows images of representative chips exposed to vehicle or candidalysin and stained for actin and nuclei. Actin positive objects and cell count (nuclei) quantification are shown in middle and bottom rows, respectively. 75 μM candidalysin significantly increased the number of actin positive objects (**Fig. 5 b**), the total actin area (**Fig. 5 c**), and the perimeter of actin objects (**Fig.5 d**) compared to vehicle. All parameters were normalized to the total cell number.

**Fig. 5.**
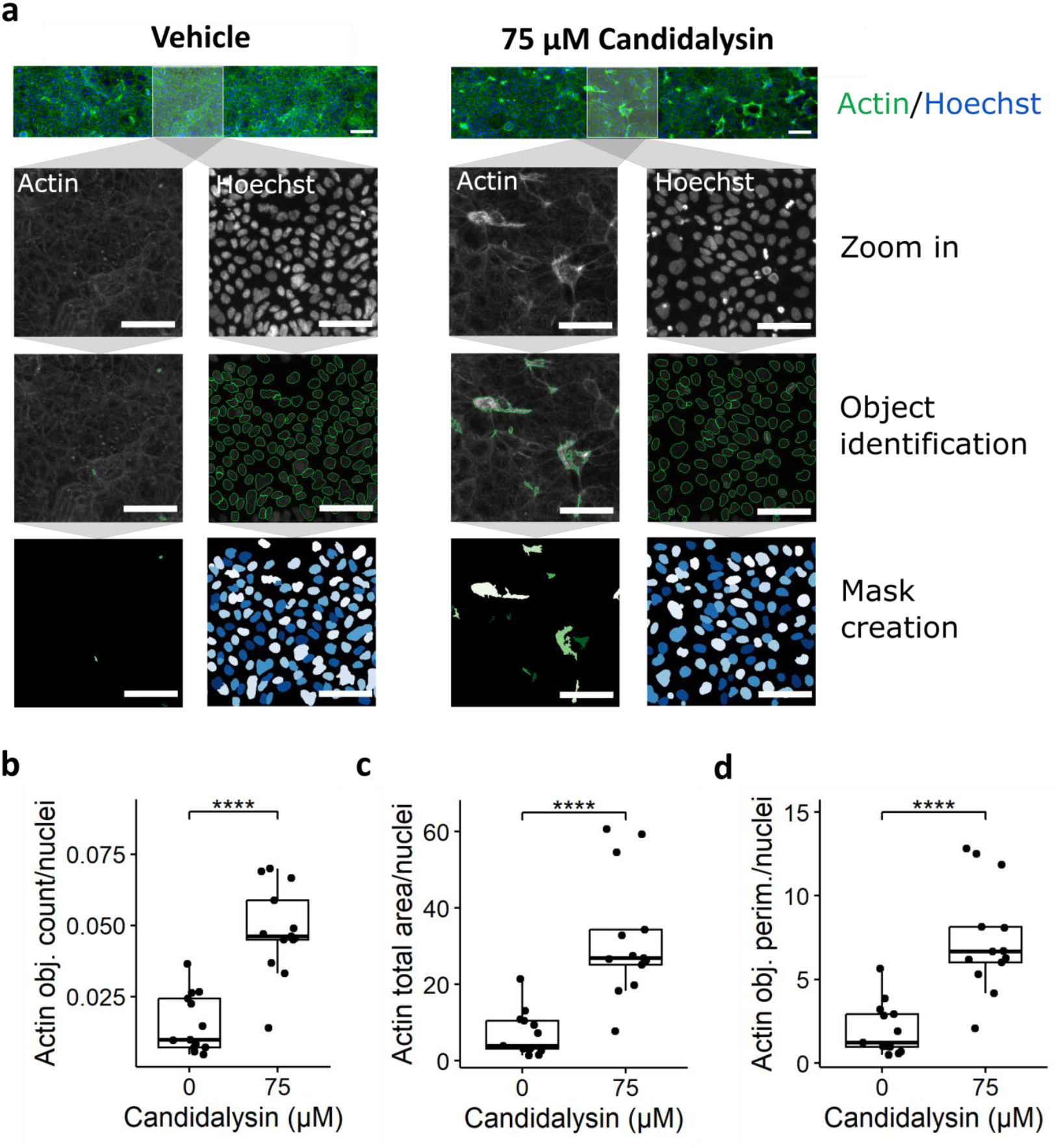
Candidalysin induces actin remodeling. Caco-2 tubules exposed to candidalysin for 2h were stained with actin and Hoechst nuclear dye. (**a**) A CellProfiler pipeline was designed to identify and quantify actin-positive and Hoechst-positive objects from confocal images. Pictures are representative of 3 independent experiments (N=3). Scalebar is 100 μM. The number (**b**), total area (**c**), and perimeter (**d**) of Actin-positive objects were normalized against the number of nuclei (Hoechst positive). Results are expressed as boxplot with individual measurements for each chip. Data was analyzed using independent T-test or Wilcoxon Rank-Sum Test (****p ≤ 0.0001, N=3, n=4-5)

### 3.3 Exposure to candidalysin induces cytotoxicity and secretion of inflammatory markers in Caco-2 tubules

To evaluate the effect of candidalysin on the cellular activation of Caco-2 tubules, the production of epithelial inflammatory mediators LL37, S100A8/A9, MCP-1, IP-10, IL-8, IL-6, IL-1beta, IL-1alpha, GM-CSF, MIP-3 alpha, and G-CSF was quantified using ELISA and Luminex technology. Caco-2 cells produced no or low amounts of these analytes in non-triggered conditions. After 2h 75 μM candidalysin, secretion of all mediators increased significantly (**Fig. 6**), except S100A8/A9, which was not detected. We used the lactate dehydrogenase (LDH) assay to measure cytotoxicity after candidalysin exposure. LDH was significantly higher in chips treated with 2h 75μM candidalysin compared to vehicle (**Fig. 6**).

**Fig. 6.**
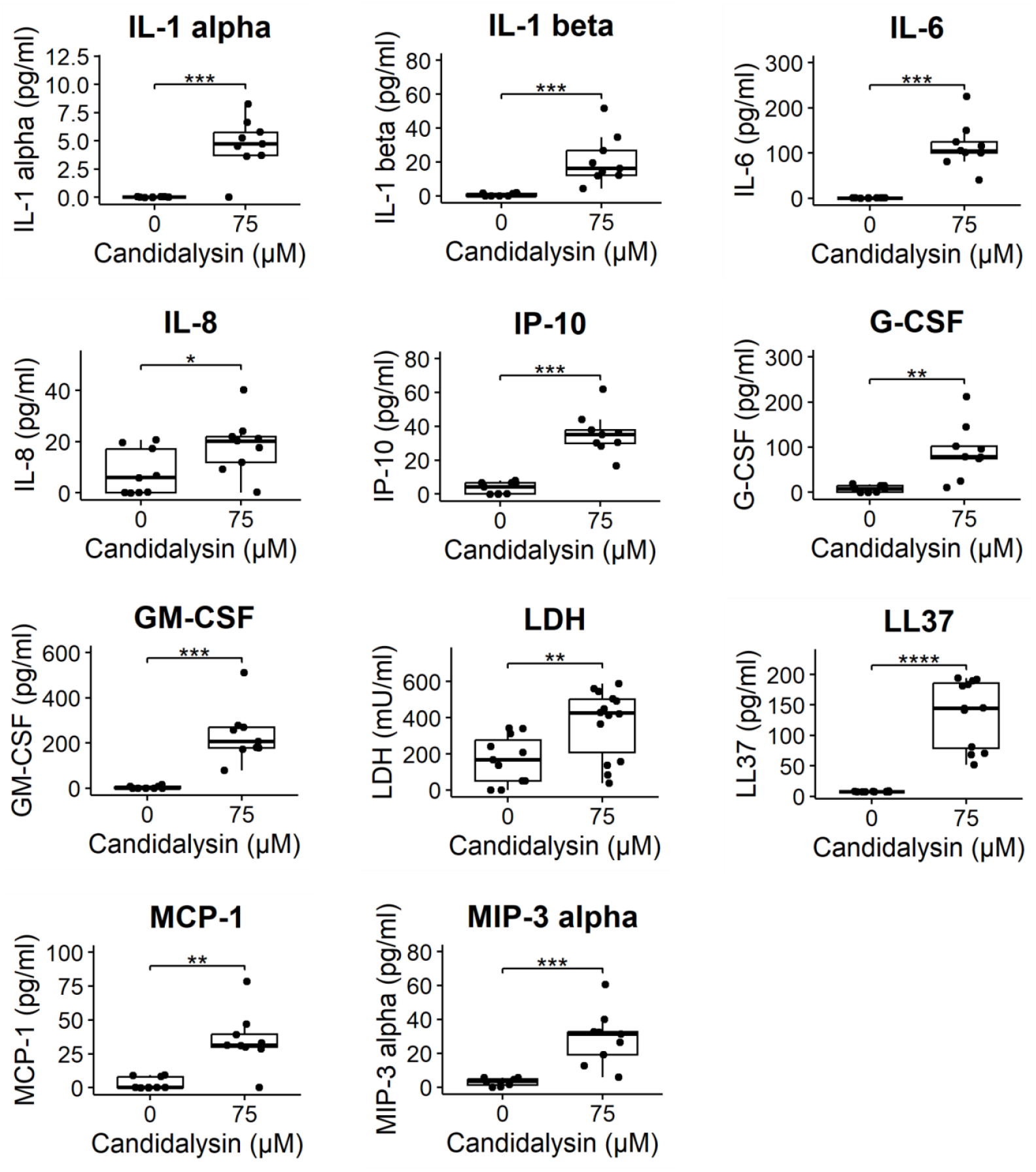
Candidalysin induces cytotoxicity and secretion of inflammatory markers in Caco-2 tubules. Candidalysin induced the secretion of MIP-3 alpha, G-CSF, GM-CSF, Il1-alpha, Il-1 beta, IL-6, IL-8, IP-10, MCP-1, LL37, and LDH in the apical side (tubule lumen). Results are expressed as boxplot with individual measurements for each chip. Data was analyzed using independent T-test or Wilcoxon Rank-Sum Test (*p ≤ 0.05; **p ≤ 0.01; ***p ≤ 0.001, ****p ≤ 0.0001, N=3, n=3-5)

### 3.4 Assessment of response to candidalysin in human colon organoids

To validate our results obtained from Caco-2 tubules and to explore future applications in a patient-derived model, we conducted a proof-of-concept experiment using human colon organoid tubules. We used the OrganoTEER to measure the barrier integrity of colon organoids following a 2-hour exposure to 0, 18, 37, and 75 μM candidalysin. Subsequently, we fixed the plates and performed DRAQ7 and actin staining (**Fig. 7 a**) and quantification. Compared to the vehicle, only 75 μM candidalysin significantly decreased TEER after the 2-hour exposure (**Fig. 7 b**). Both 37 and 75 μM candidalysin increased permeability to DRAQ7 compared to the vehicle (**Fig. 7 c**). Furthermore, 75 μM candidalysin led to significant actin remodeling, as evidenced by increases in the number (**Fig. 7 d**), total area (**Fig. 7 e**), and perimeter of actin objects (**Fig. 7 f**).

**Fig. 7.**
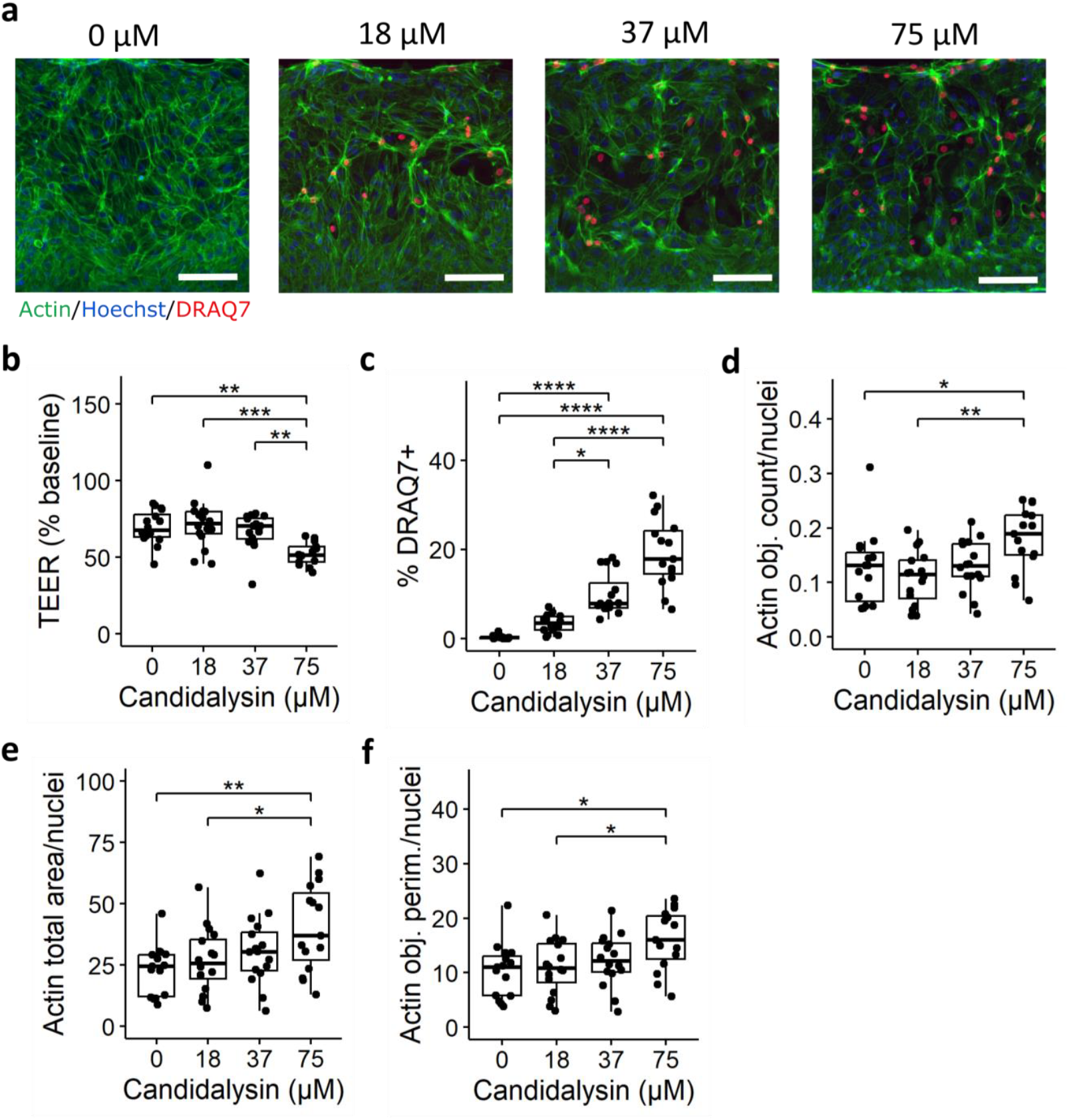
Candidalysin exposures in human colon organoid tubules. (**a**) colon organoid tubules exposed to 0, 18, 37, and 75 μM candidalysin for 2h were stained with actin (green), DRAQ7 (red), and Hoechst (blue). Representative images are max projections taken at the bottom of the tubules. Scale bar is 100 μM. (**b**) TEER measurement after candidalysin exposure. Each chip is normalized to its value at baseline (before exposure). (**c**) The number (**d**), total area (**e**), and perimeter (**f**) of actin-positive objects were normalized against the number of nuclei (Hoechst positive). Results are expressed as boxplot with individual measurements for each chip. Data was analyzed using Kruskal Wallis test with Dunn’s post-hoc test or one-way ANOVA with Tukey’s HSD post-hoc test (*p ≤ 0.05; **p ≤ 0.01; ***p ≤ 0.001, ****p ≤ 0.0001, N=3, n=3-6)

## 4 Discussion

This study aimed to evaluate the phenotypic response of IECs to candidalysin, focusing on barrier integrity and inflammation. Given that candidalysin forms pores and damages the epithelium[13, 14], we hypothesized it would impair barrier integrity and increase permeability in our model. We also expected to detect toxicity and inflammation mediators, as described in studies using oral epithelial cells[17, 19].

To test these hypotheses, we used the OrganoPlate technology, allowing the culture of up to 64 perfused intestinal Caco-2 tubules in a membrane-free system[24]. This platform allowed the rapid assessment of the barrier integrity of the gut tubules[25], along with easy imaging and sampling of luminal medium to detect inflammatory markers. We found that candidalysin impaired barrier integrity of Caco-2 tubules (measured by TEER and barrier integrity assay), increased permeability (measured by DRAQ7 staining), and induced actin remodeling and secretion of inflammatory mediators. We then validated our results in a more complex model using patient-derived organoid tubules exposed to candidalysin, measuring TEER, DRAQ7 permeability, and actin remodeling.

Disruption of barrier integrity is a hallmark of mucosal inflammation. Using a combination of readouts, we observed that candidalysin impaired barrier integrity and increased permeability. In Caco-2 tubules, only TEER detected significant differences at all timepoints after 37 μM candidalysin exposure, suggesting that TEER is more sensitive than barrier integrity assay[24, 25]. In colon organoid tubules, TEER was less sensitive than in Caco-2, with only 75 μM of candidalysin showing a significant decrease after 2-hour exposure. This reduced sensitivity may be due to the increased complexity of colon organoids compared to Caco-2 and could be influenced by the washing step before exposure, as indicated by decreased TEER in the vehicle control.

While several studies reported loss of barrier integrity after infection with *C. albicans* in IECs[7, 10, 27] and other epithelial cell type[28, 29], to date, only one study used exogenous candidalysin in IECs. Contrary to our observations, they did not observe a decrease in TEER or an increase in 4-kDa dextran permeability after 24 hours of 70 μM candidalysin exposure using the Caco-2 subclone C2BBe1 in Transwells[10]. This discrepancy likely arises from model differences (dynamic gut-on-chip vs. static Transwells), exposure duration (acute 2-hour vs. prolonged 24-hour), cell line sensitivity, and measurement sensitivity (more responsive TEER in the gut-on-chip[25]). These differences highlight how model choices impact findings, suggesting the gut-on-chip system could better capture acute barrier disruption, influencing interpretations of candidalysin’s role in epithelial damage.

The actin cytoskeleton mediates disruption of mucosal barriers in inflamed tissues[30]. We therefore looked at the actin network after candidalysin exposure and observed remodeling in Caco-2 and organoids, confirming results observed in oral epithelial cells[21]. Notably, in colon organoids, actin clumps were found to co-localize with Draq7-positive nuclei, suggesting a localized effect of candidalysin in these areas.

Similarly to observations in other cell types[12, 17–19], we observed the induction of several candidalysin-induced proteins in Caco-2. Along with the release of LL-37, G-CSF, GM-CSF, IL-6, IL-1a, IL-1b, MCP-1, and IL-8, we also observed candidalysin-induced release of IP-10 and MIP-3 in our cultures. Notably, S100A8/A9 release was not induced by candidalysin in our system, in line with findings in TR146 cells[17]. The secretion of LL-37, an antimicrobial peptide, is induced by candidalysin, highlighting its role in the host defense mechanism. However, further experiments with additional stimuli are needed to determine the specificity of this response. Future studies could address this by testing other toxins or stimuli to compare their effects on LL-37 release and clarify whether this response is specific to candidalysin. These observations support the notion that the epithelial barrier is not just a physical barrier but actively participates in the host’s early response to pathogens[31, 32].

A limitation of our study is the use of exogenous candidalysin. While it allows for the evaluation of its direct effect on IECs and provides insights into the early stages of *C. albicans* infection, it is important to note that the delivery of candidalysin to host cell membranes is critical[33] and is not recapitulated in our system. Additionally, our model does not address direct fungal-host interactions, such as the use of host zinc by the yeast to promote translocation and virulence[23]. Another limitation is the lack of immune components. Gut-on-chip models of *C. albicans* infection including living yeast and immune cells have been reported[34] but still present technical challenges to be scaled up for testing potential therapies[35]. Moreover, advanced gut-on-chip models that enable the study of immune migration have been described[36, 37], and their adaptation to investigate candidiasis pathogenicity could be a promising direction. Another point to consider is the interactions between *C. albicans* and the rest of the microbiota, as it has been shown that candidalysin can inhibit bacterial species[38].

In this study, protein detection was not measured in the organoid tubules due to the exploratory nature of this approach. Future experiments should incorporate this approach to leverage the more physiologically relevant environment of organoids, potentially providing more accurate insights into protein expression responses. Furthermore, the response to candidalysin might be donor-specific, highlighting the need to test in more donors to capture the variability in responses. This research represents a starting point for understanding the physiological responses to candidalysin and underscores the necessity of screening more donors while examining various response angles.

In summary, our gut-on-chip model effectively evaluated the phenotypic response to candidalysin in intestinal epithelial cells, providing insights into the toxin’s mechanisms. This system’s ability to multiplex readouts offers a valuable tool for screening potential therapeutic inhibitors and modulators of candidalysin-induced responses. It could be used to screen for candidalysin inhibitors, such as nanobodies neutralizing candidalysin[39], or modulators of candidalysin-induced responses. Furthermore, this model can be adapted to study the effects of candidalysins produced by other non-*albicans Candida* species, which are also prevalent fungal pathogens[19]. It could also be adapted to study the phenotypic response of other molecules like other toxins[40], micro-organisms, and microbial metabolites. Lastly, the use of patient-derived organoids allows for personalized modeling and the use of cells coming from patients with candidiasis or other diseases. Overall, our findings highlight the potential of our platform to study the effects of various virulence factors produced by different microorganisms and microbial communities. This paves the way for developing targeted therapeutic strategies that modulate these factors or the host responses they trigger.

## 5 Conclusion

This study demonstrates the utility of a gut-on-chip platform for evaluating the toxicological effects of candidalysin on intestinal epithelial cells. By integrating dynamic, membrane-free models and patient-derived organoids, we have characterized candidalysin-induced epithelial damage and inflammatory responses in a physiologically relevant system. These findings not only enhance our understanding of *Candida albicans* pathogenesis but also highlight the potential of this platform for toxicology screening and therapeutic development. Future applications include expanding the model to incorporate immune components and studying the effects of other microbial toxins, paving the way for personalized approaches to managing fungal infections.

## 7 Author Contributions

M.M. designed and performed the experiments, analyzed the data, and wrote the manuscript. KQ provided guidance and revision. All authors have read and agreed to the published version of the manuscript.

## 9 Acknowledgments

The authors would like to thank Bernhard Hube, Mark Gresnigt, and Selene Mogavero of the Hans-Knöll institute in Jena, Germany for their insights on candidalysin usage and for helpful discussions on this work. We thank Frederik Schavemaker at MIMETAS for designing some of the illustrations. We also thank our colleagues at MIMETAS and in the FunHoMic consortium for the many fruitful discussions.

## 12 Data Availability Statement

The datasets generated during this study are available from the corresponding authors on reasonable request.

## 13 Ethical statement

The organoid models used in this study (OrganoReady Colon Organoid) were obtained as ‘ready-to-use’ products from MIMETAS B.V. The cells used to generate these models were sourced by MIMETAS from a third-party tissue provider. This provider has confirmed that all human tissue samples were collected in compliance with applicable laws and ethical guidelines, including (but not limited to) the Declaration of Helsinki and standards set by organizations such as the National Bioethics Advisory Committee, American Medical Association, and Human Tissue Authority (HTA). Informed consent was obtained from all donors by the tissue provider, and ethical oversight was managed by the tissue provider and MIMETAS. No additional ethical approval was required for this study.

