## Supplementary information for "Breaking barriers: the fungal toxin candidalysin disrupts epithelial integrity and induces inflammation in a gut-on-chip model"

### Supplementary Materials

Table 1. Table of reagents

| Reagent | Supplier | Reference number | Note |
|---|---|---|---|
| Fluorescein sodium salt | Sigma-Aldrich | F6377-100G | Stock: 1mg/ml in water<br>Working: 10 µg/ml |
| TRITC-Dextran 4.4 kDa | Sigma-Aldrich | T1037 | Stock: 25mg/mL in water<br>Working: 500 µg/ml |
| ActinGreen™ 488 ReadyProbes™ Reagent | Sigma-Aldrich | R37110 | 2 drops/ml |
| NucBlue™ Live ReadyProbes™ Reagent | Sigma-Aldrich | R37605 | 2 drops/ml |
| DRAQ7™ Dye | Biostatus | DR71000 | Working: 3 µM |
| Triton 100X | Sigma-Aldrich | T8787 | |
| Formaldehyde | Sigma-Aldrich | 252549 | |
| HBSS | ThermoFisher | 14025-092 | |
| PBS | Gibco | 70013-016 | Diluted 10X in milliQ |
| EMEM | ATCC | 30-2003 | Caco-2 medium:<br>440mL EMEM<br>50mL FBS<br>5mL P/S<br>5mL NEAA |
| FBS | Gibco | 16140-071 | |
| P/S | Gibco | 15140-122 | |
| MEN NEAA | Gibco | 11140-050 | |

| | | | |
|---|---|---|---|
| OrganoReady Colon Caco-2 3-lane 40 | MIMETAS | MI-OR-CC-01 | Use caco-2 medium |
| OrganoReady Colon Organoid 3-lane 64 | MIMETAS | MI-OR-CORG-02 | Includes medium:<br>• OrganoMedium Colon Organoid-ARM Apical Recovery Medium<br>• OrganoMedium Colon Organoid-BRM Basolateral Recovery Medium<br>• OrganoMedium Colon Organoid-ACM Apical Culture Medium<br>• OrganoMedium Colon Organoid-BCM Basolateral Culture Medium |
